# Early embryogenesis in CHDFIDD mouse model reveals facial clefts and altered craniofacial neurogenesis

**DOI:** 10.1101/2023.04.23.537955

**Authors:** M Hampl, N Jandova, D Luskova, M Novakova, J Prochazka, J Kohoutek, M Buchtova

## Abstract

Congenital heart defects, facial dysmorphism and intellectual development disorder (CHDFIDD) is associated with mutations in *CDK13* gene which encodes a transcription regulating Cyclin-dependent kinase 13 (CDK13). Here we analyzed early embryonic stages of CHDFIDD mouse models with hypomorphic mutation in *Cdk13* gene with very similar phenotypic manifestations plus cleft lip/palate and knockout of *Cdk13* which exhibits robust phenotype with midfacial cleft. *Cdk13* is strongly expressed in the mouse embryonic craniofacial structures, namely in the forebrain, nasal epithelium and maxillary mesenchyme. *In vitro,* CDK13 protein is located not only in nuclear region but also in the cellular protrusions in cultured mesenchymal cells and cells isolated from dorsal root ganglia. In *Cdk13*-deficient embryos, we found hypoplastic branches of the trigeminal nerve including maxillary branch and additionally we detected significant gene expression changes of molecules involved in neurogenesis (*Mef2c*, *Pou4f1*, *Sod1*, *Cdk5rap2*, *Nrcam*) within the developing palatal shelves. Key palate-associated molecules (*Msx1* and *Meis2*) were downregulated during early craniofacial development in mutant embryos. These results demonstrate the role of CDK13 in regulation of facial morphogenesis and also growth of craniofacial peripheral nerves.

## INTRODUCTION

CDK13 is one of the transcriptional kinases, which regulates transcription via phosphorylation of the RNA Polymerase II and control alternative splicing (Bartkowiak et al., 2010, Blazek et al., 2011). Recently, a few case studies have described a variety of developmental defects in human patients with *CDK13* mutation (Hamilton and Suri, 2019). These patients exhibit delayed development, intellectual disorders, heart and kidney defects and craniofacial deformities. These features have been recognized as Congenital heart defects, facial dysmorphism and intellectual development disorder (CHDFIDD) syndrome. In addition to the most common defects, patients suffer from brain defects, autism, seizures, limb defects, defects in skeletogenesis and other abnormalities (Hamilton and Suri, 2019). To simulate defects presented in patients with *CDK13* mutations, we developed mouse models exhibiting similar phenotype to human (Nováková et al., 2019). We observed that *Cdk13*-deficiency in mouse model causes embryonic lethality, delayed development, heart and brain abnormalities (Nováková et al., 2019). Here, we focus on the alteration of developmental processes in craniofacial structures in the *Cdk13*-hypomorph (*Cdk13^tm1a/tm1a^*) and knock-out (*Cdk13^tm1d/tm1d^*) mouse embryos with aim to uncover cellular mechanisms behind observed developmental abnormalities. Interestingly, previous, *in vitro* approaches uncovered contribution of *Cdk13* mutation to alteration of neuronal differentiation and neurite outgrowth through regulation of the CDK5 pathway (Chen et al., 2014). These findings are in agreement with observations of neurodevelopmental disorders in human patients bearing *CDK13* mutation (Trinh et al., 2019). Connection between insufficient growth of craniofacial nerves leading to development of various craniofacial clefts has been already described (Rizos et al., 1998, Gao et al., 2013). Based on this evidence, we hypothesize that *Cdk13*-deficiency in mouse embryos could also lead to altered growth of craniofacial nerves which could lead to development of severe facial clefts which we observed in both hypomorph and knockout embryos (Nováková et al., 2019). Here, we aim to evaluate possible alterations of individual branches of the trigeminal nerve and detect expression changes of neurogenesis specific genes and craniofacial structures characteristic molecules to find possible signaling pathways and cellular processes, which can lead to defects in morphogenesis of the craniofacial structures.

## RESULTS

### *Cdk13*-deficiency causes craniofacial defects including severe facial clefting

In our previous study, we observed craniofacial developmental defects caused by *Cdk13*-deficiency (Nováková et al., 2019). Here we analyzed all critical stages of *Cdk13^tm1a/tm1a^* and *Cdk13^tmda/tm1d^*embryos which exhibited morphological defects of craniofacial structures including severe clefting (Fig.1Q). Protruding neural tube to the developing face and missing midline facial parts were detected in E11.5 *Cdk13^tm1d/tm1d^* embryos (Fig.1A′′). In E12.5, we found cleft lip in *Cdk13^tm1a/tm1a^* embryos (Fig.1B′,H′), missing midline structures in *Cdk13^tm1d/tm1d^* embryos resulting in midline cleft (Fig.1B′′,H′′), and delayed palatal shelves (PSs) development (Fig.1É, K′,K′′) with cleft nasal septum (Fig.1K′,K′′) in both mutant genotypes. Later at E13.5 and E14.5, *Cdk13^tm1a/tm1a^* embryos showed thinner lips with wider distance between (Fig.1F′,Í,G′J′) and delayed development of the PSs (Fig.1F′,Ĺ,G′,M′), while *Cdk13^tm1d/tm1d^* embryos showed massive midfacial cleft (Fig.1Ć′,Í′,D′′,J′′) and persisting nasal septum cleft (Fig.1Ĺ′,M′′). Less severe defects were detected in the caudal palate area, where the most prominent were underdeveloped PSs, especially in *Cdk13^tm1d/tm1d^*embryos (Fig.S1E-G). In E16.5 *Cdk13^tm1a/tm1a^* embryos (lethality stage), cleft palate still persisted with insufficient horizontalization in the rostral area, but not in the caudal region, animals still did not exhibit fused lips in the midline (Fig.S1B-B′′,D,D′).

**Figure 1.**
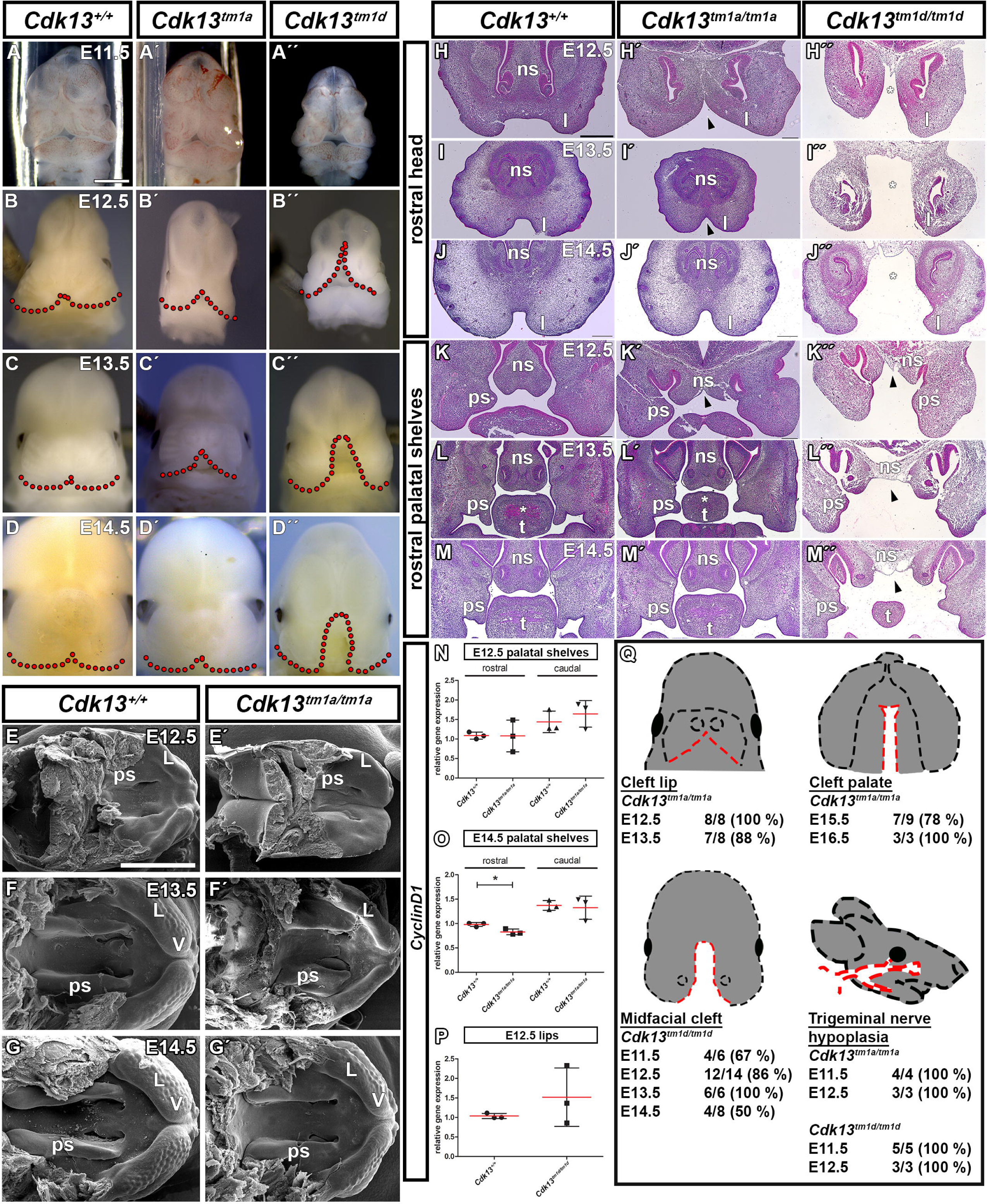
*Cdk13-*deficient embryos display severe craniofacial clefting. **(A – D′)** External craniofacial phenotype of the E11.5 - E14.5 embryos. Dotted lines highlight shape of the upper lip on frontal views. **(É - G′)** Palatal view of the E12.5 – E14.5 embryos using scanning electron microscope showing differences in morphology of the palatal shelves, forming lips and vibrissae. **(H – M′′)** Hematoxylin-Eosin staining of the E12.5 – E14.5 frontal head sections. **(H – J′′)** Arrowheads point to cleft lip in hypomorph (*Cdk13^tm1a^*) embryos and asterisks highlight midfacial cleft in KO (*Cdk13^tm1d^*) embryos. **(K – M′′)** Arrowheads point to cleft nasal septum in both hypomorph and KO embryos. The palatal shelves development in rostral region is altered in both genotypes compared to WT embryos. Hypoplastic muscles in tongue are marked by asterisks. (**N - P)** qPCR quantification of changes in proliferation shown as *CyclinD1* gene expression in the rostral **(N)** and caudal **(O)** palatal shelves of the E12.5 and E14.5 hypomorph embryos and in lips of the E12.5 KO **(P)** embryos. t - test; **0.001 < p < 0.01; *p < 0.05. **(Q)** Table displays frequency of severe craniofacial phenotypes (cleft lip, cleft palate, midfacial cleft, hypoplasia of the trigeminal nerve) within individual genotypes and embryonic developmental stages. Red dashed lines contour defective structures. L – lip; ns – nasal septum; ps – palatal shelf; t – tongue; v - vibrisae. Scale bars: macroscopic pictures - 1 mm, Hematoxylin-Eosin stained sections – 500 μm, SEM pictures – 1 mm.

Although the PSs in the *Cdk13*-deficient animals were generally smaller and exhibited different shape, there were no significant changes detected in proliferation (*cyclinD1*) in the PSs of *Cdk13^tm1a/tm1a^*and lips of *Cdk13^tm1d/tm1d^* embryos (Fig.1N-P). Additional immunostaining of Ki-67 and TUNEL assay confirmed no significant changes in proliferation and apoptosis in the developing palatal shelves of *Cdk13^tm1a/tm1a^* embryos (Fig.S4A-C).

### Expression of *Cdk13* at mRNA level is dispersed through developing facial regions

As we observed distinct changes in facial and palatal morphogenesis in *Cdk13*-deficient embryos, we asked if there is localized expression of the *Cdk13* mRNA to certain areas during craniofacial development. In the earlier developmental stages (E11-E14), *Cdk13* was expressed in both rostral and caudal areas of the developing PSs with more dense *Cdk13* signal detected in the rostral region (Fig.2B′′;Fig.S2F,G) compared to caudal areas (Fig.2B′;Fig.S2F,G). In the developing snout, intensity of the *Cdk13* signal was similar in the surface epithelium and mesenchyme. Visibly stronger signal was detected in the E11 forebrain (Fig.2C) and nasal epithelium (Fig.2C′,Ć′) and in the E12 maxillary mesenchyme (Fig.2D). Changes emerged in older embryos (E15-E16), when we observed strong expression of *Cdk13* in the palatal epithelium and adjacent mesenchyme, especially in the developing palatal rugae, both in the rostral and caudal (Fig.S2H,I) regions. Quantification of *Cdk13* expression by qPCR revealed its decreasing level in the palatal area through the development (Fig.2E) and increasing level in the maxillary, mandibular and frontonasal prominences (Fig.2F).

**Figure 2.**
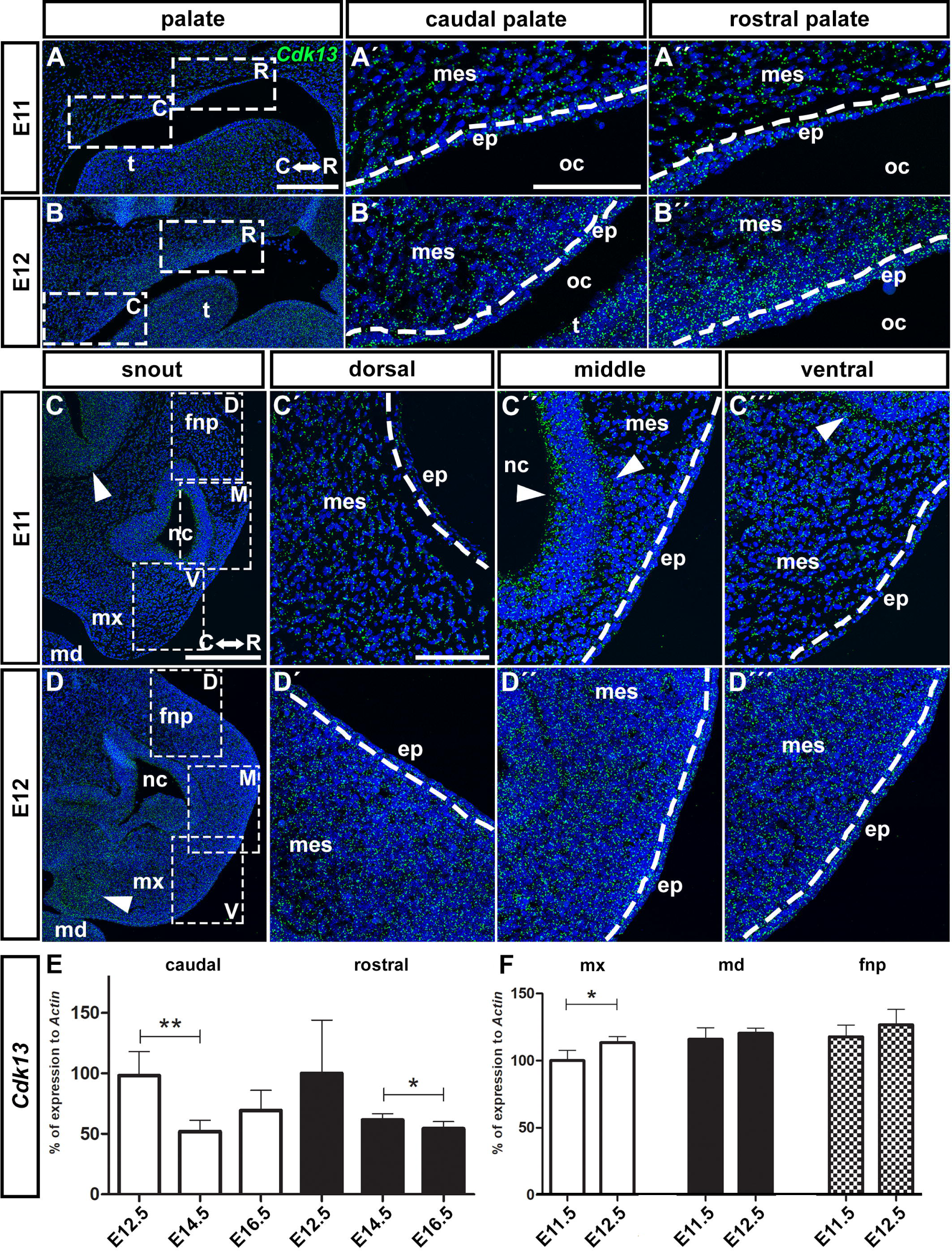
Physiological gene expression of the *Cdk13* in the developing palatal tissues and snout on sagittal sections detected by RNAScope and its quantification by qPCR. **(A–B)** Lower power magnification of the *Cdk13* (green) gene expression in the developing palatal tissues at E11 and E12 stages of WT animals. White dashed line rectangles highlight regions used for higher power pictures in caudal (C; **A′-B′)** and rostral (R; **A′′-B′′)** regions. **(C–D)** Lower power magnification of the *Cdk13* gene expression in the developing snout at E11 and E12 stages of WT animals. White dashed line rectangles highlight regions used for higher power pictures in dorsal (D; **Ć, D′)**, middle (M; **Ć′,D′′)**, and ventral (V; **Ć′′,D′′′)** areas. Arrowheads point to dense signal in the developing forebrain **(C)**, nasal epithelium **(Ć′, Ć′′)** at E11 and in the developing mx **(D)** at E12. White dashed lines separate mesenchyme and epithelium. Nuclei are counterstained with DAPI. **(E)** Quantification of the *Cdk13* gene expression by qPCR in the E12.5, E14.5 and E16.5 palatal shelves. White columns display expression in the caudal palate and black columns expression in the rostral palate. **(F)** Quantification of the *Cdk13* gene expression by qPCR in the E11.5 and E12.5 facial prominences: white columns display expression in the mx, black in the md and patterned in the fnp. Unpaired two-tailed Student t - test; **0.001 < p < 0.01; *p < 0.05. c – caudal; ep – epithelium; fnp – frontonasal prominence; mes – mesenchyme; md – mandibular prominence; mx – maxillary prominence; nc – nasal cavity; oc – oral cavity; r – rostral; t – tongue. Scale bars: lower power – 200 μm, higher power – 100 μm.

To uncover possible differences of *Cdk13* expression through PSs in labio-lingual direction, we evaluated distribution of *Cdk13* on frontal sections. In E12-E14 embryos, *Cdk13* signal was spread evenly within the palatal mesenchyme and epithelium of the PSs (Fig.S2A-C) and later in E15 and E16 embryos, we detected enriched expression of *Cdk13* in the palatal mesenchyme close to region of fusion in the craniofacial midline (Fig.S2D,E).

### CDK13 protein is localized in the cellular outgrowths as well as long neural processes

Next, we asked how CDK13 protein is distributed in cells *in vitro*. We analyzed its expression in cultured MEF cells (mouse embryonic fibroblasts) and cells derived from DRGs (dorsal root ganglia) of adults as these cell types are typical by formation of long cytoplasmic processes.

In MEF cells, the CDK13 was localized in nuclear area and enriched in cytoplasm around nucleus as predicted by its function as a regulator of transcription (Fig.3A). However, CDK13 protein was also detected in the cellular outgrowths (Fig.3A′-A′′′) and it was enriched especially in their most apical tips where it was located together with actin filaments (Fig.3A′). In cells isolated from DRGs, CDK13 was detected in the nuclear area with the strongest expression detected in the cytoplasm closely adjacent to nuclei (Fig.3B). Similarly to MEF cultures, we detected CDK13 in long neural processes along the neurofilaments, which were forming the outgrowths (Fig.3B′,B′′). This specific cellular localization indicates a potentially new role of CDK13 protein, in addition to regulation of RNA Polymerase II activity in nucleus, as involved in the interaction with cytoskeletal proteins and contribution to cell protrusions establishment.

**Figure 3.**
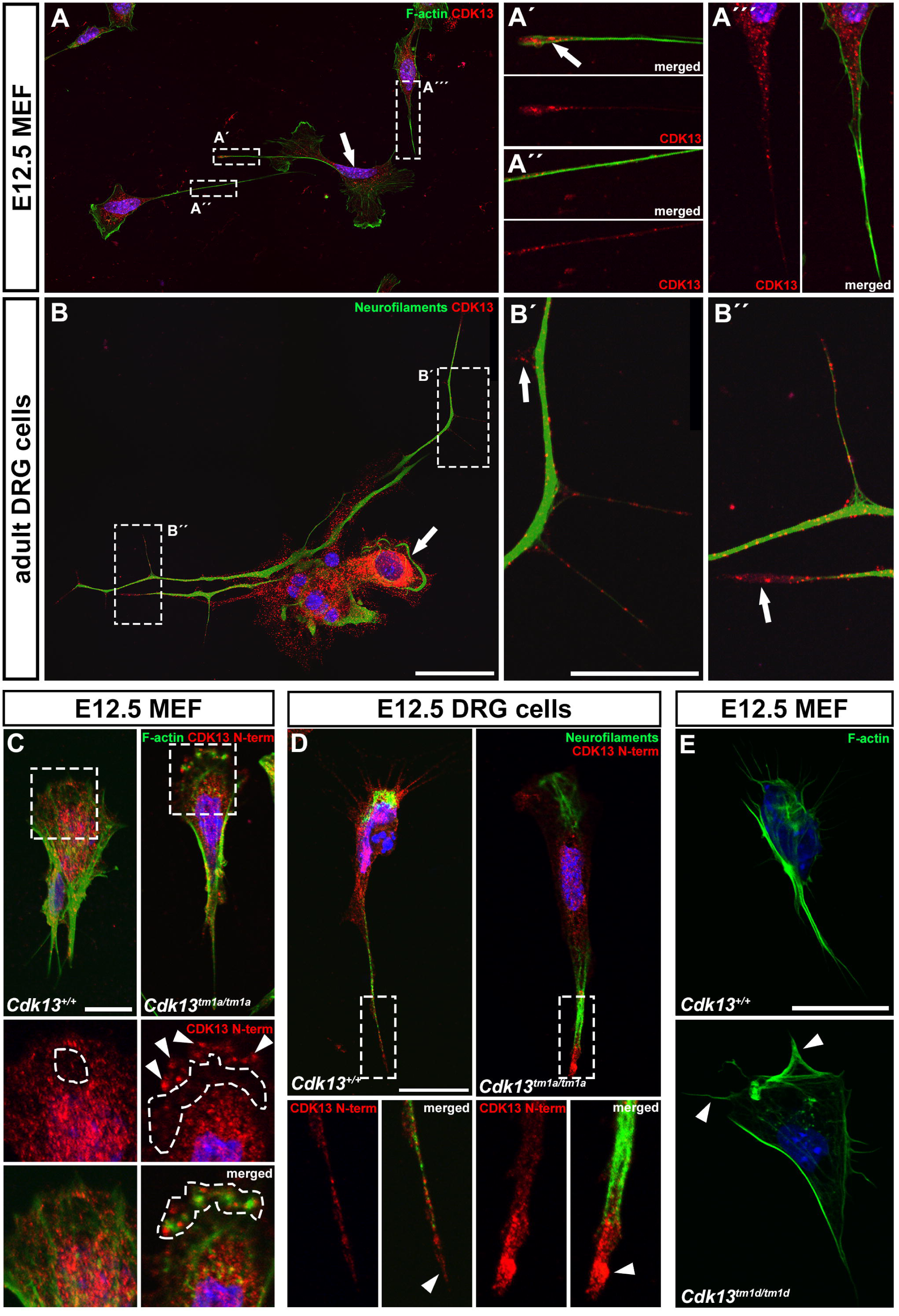
*In vitro* immunocytochemical detection of the CDK13 in different cell types. **(A)** Lower power picture of CDK13 (red) and F-actin (green) expression in mouse embryonic fibroblasts (MEF) isolated from E12.5 embryonic bodies. Arrow points to nuclear area with CDK13 expression. Dashed line rectangles highlight details of cellular outgrowths. **(A′**-**A′′′)** Details of CDK13 expression in the forming cellular processes. Arrow points to apical area with enriched CDK13 expression **(A′)**. **(B)** Lower power picture of CDK13 (red) expression and neurofilaments (green, anti-2H3) in primary cells isolated from dorsal root ganglia of adult mice. Arrow points to nuclear area with strong CDK13 expression. Dashed line rectangles highlight details on neurite outgrowths. **(B′**, **B′′)** Details of CDK13 expression in the forming neurite outgrowths. Arrows point to areas with CDK13 expression prior to neurofilaments formation. Nuclei are counterstained with DAPI. **(C)** Detection of the truncated form of CDK13 (red) in E12.5 *Cdk13^tm1a/tm1a^* MEF cells using anti N-terminal end CDK13 antibody. Dashed line rectangles highlight details on cellular outgrowths. Detailed pictures focus on aggregates of the truncated CDK13 in the cellular protrusions (arrowheads), CDK13-negative area (dashed line region, red) and colocalization of the truncated CDK13 with F-actin (green) deposits (dashed line region, merged). **(D)** Detection of the truncated form of CDK13 (red) in E12.5 *Cdk13^tm1a/tm1a^* DRG cells using anti N-terminal end CDK13 antibody colocalized with neurofilaments (green, anti-2H3). Dashed line rectangles highlight details on long cellular outgrowths. Detailed pictures focus on accumulation of the truncated CDK13 in distal part of the cellular outgrowth (arrowhead, merged). **(E)** Detection of the cellular protrusions by F-actin antibody (green) in E12.5 *Cdk13^tm1d/tm1d^*MEF cells. Less and thicker cellular protrusions (arrowheads) produced by cells isolated from mutant embryos. Scale bars: (A-B′′) - lower power – 50 μm, higher power – 20 μm; (C) – 10 μm; (D) – 20 μm; (E) - 20 μm.

Hypomorphic mutation causes production of the truncated form of CDK13 with preserved N-terminal domain (Nováková et al., 2019). This form of CDK13 aggregated in the cellular processes together with deposits of F-actin in E12.5 *Cdk13^tm1a/tm1a^*MEF (Fig.3C). Similarly, aggregates of the truncated CDK13 were located in wide protrusions of cells isolated from E12.5 *Cdk13^tm1a/tm1a^* DRGs (Fig.3D). Additionally, thick and less amount of cellular protrusions was also observed in MEF isolated from E12.5 *Cdk13^tm1d/tm1d^* (Fig.3E; Fig.S3A,B).

### *Cdk13*-deficiency results in hypoplastic craniofacial nerves

Development of the craniofacial region is closely associated with development of the craniofacial nerves. The largest area is innervated by trigeminal nerve with three main branches: maxillary, mandibular and ophthalmic (Higashiyama and Kuratani, 2014). We detected complex spatial *Cdk13* expression in the developing maxillary nerve (MxN) in cells ensheathing bundles of nerve fibers, localized in their nuclei and also in the cytoplasmic projections (Fig.S3C-G′). As the CDK13 was demonstrated to regulate neurite outgrowth (Chen et al., 2014), we hypothesized that alteration of craniofacial nerves growth could occur during early craniofacial development in *Cdk13*-deficient animals.

First, we visualized outgrowth of cranial nerves by whole mount immunohistochemistry analysis of neurofilaments in *Cdk13^tm1a/tm1a^*and *Cdk13^tm1d/tm1d^* embryos (Fig.4B-Ć) to uncover possible alterations of general morphology of trigeminal nerve and associated nerves. Alteration of several craniofacial nerves outgrowth was detected in *Cdk13^tm1a/tm1a^* and *Cdk13^tm1d/tm1d^* embryos (Fig.4B-Ć) with obvious hypoplasia of maxillary (MxN), mandibular (MdN) and ophthalmic (OpN) nerves, which were reduced in length. These nerves originate from the trigeminal ganglion (TG), which we found to be also abridged in the mutant embryos (Fig.4B,Ć′). Only frontal nerve (frN) was not found to be morphologically altered (Fig.4B′,Ć). This confirms that *Cdk13*-deficiency negatively affects neurogenesis during embryonic development and demonstrates that CDK13 is involved in the craniofacial nerve outgrowth.

**Figure 4.**
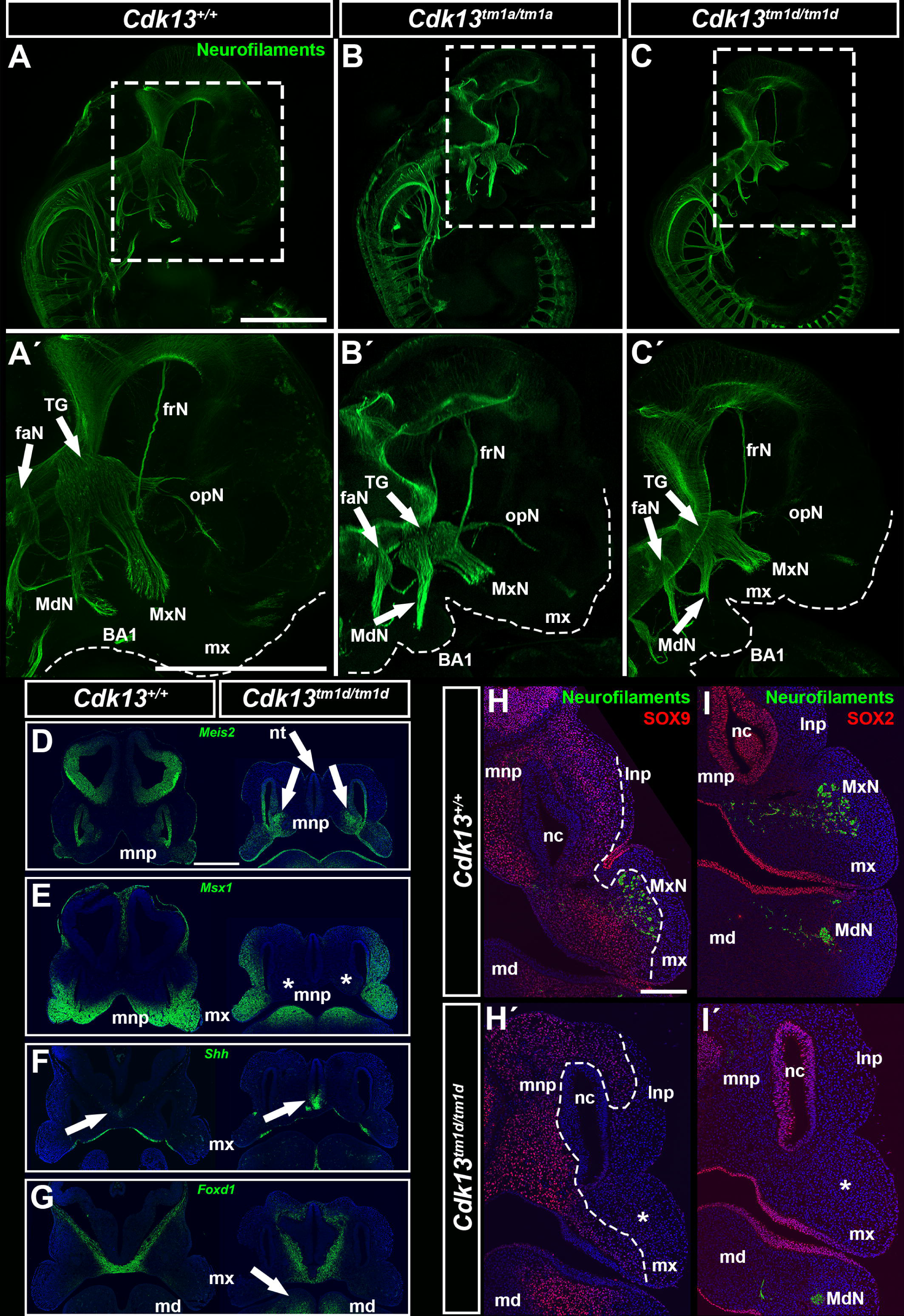
Immunohistochemical detection of craniofacial nerves on whole mount embryos and on sections supplemented with RNAScope *in situ* gene expression on sections of the *Cdk13*-deficient embryos. **(A-C)** Development of neurons detected using neurofilaments antibody (green, anti-2H3) on the E11.5 WT, *Cdk13^tm1a/tm1a^* and *Cdk13^tm1d/tm1d^* embryos. Dashed line rectangle highlights detail of craniofacial region. **(A′-Ć)** Higher power pictures focused on craniofacial nerves in E11.5 embryos. Detail displays the basis of the facial nerve, and divisions of the trigeminal nerve – mandibular, maxillary and ophthalmic nerves, including frontal branch nerve of the ophthalmic nerve. Dashed line highlights edges of the maxillary prominences and branchial arch 1. Note underdeveloped and hypoplastic all three divisions of the TG nerve in *Cdk13^tm1a/tm1a^* **(B′)** and especially in *Cdk13^tm1d/tm1d^***(Ć)** embryos. **(D-G)** RNAScope detection of the *Meis2*, *Msx1*, *Shh* and *Foxd1* gene expression on the frontal sections of E11.5 embryos. Gene expression was detected in the maxillary, mandibular and in the lateral and medial nasal prominences. **(H-Í)** Immunohistochemical detection of neurofilaments (anti-2H3), SOX9 and SOX2 on frontal sections of E11.5 embryos. Missing maxillary nerves in *Cdk13*-deficient embryos marked by asterisks **(H′,Í)**. White dashed lines label areas of the SOX9 expression **(H,H′)**. Scale bars: whole mount - 1 mm; RNAScope – 500 μm; IHC sections – 20 μm. BA1 – 1. branchial arch; faN – frontal and acoustic nerves; frN – frontal branch of the ophthalmic nerve; lnp – lateral nasal prominence; MdN – mandibular branch of the trigeminal nerve; md – mandibular prominence; mnp – medial nasal prominence; mx – maxillary prominence; MxP – maxillary branch of the trigeminal nerve; nc – nasal cavity; nt – neural tube; opN – ophthalmic branch of the trigeminal nerve; ot – otic vesicle; TG – trigeminal ganglion.

Similar to whole mount IHC, changes of nerve protrusions were even more obvious on transversal sections of E11.5 *Cdk13^tm1d/tm1d^*embryos. We observed in the rostral region missing MxN in the developing maxillary prominence (Fig.4H′,Í), hypoplastic MnN in the mandibular prominence (Fig.4Í) and reduced size of the TGs (Fig.S3H,I) confirming defective nerve growth in *Cdk13*-deficient animals.

This effect of *Cdk13*-deficiency on peripheral nerves development was additionally tested by chemical inhibition of CDK13 together with CDK12 (THZ531 inhibitor, there is no CDK13-specific inhibitor, thus we used the THZ531 compound inhibitor) in functional *ex vivo* cultivation experiment using trigeminal ganglia explants. TGs were dissected from E12 WT embryos and cultured supplemented with THZ531 inhibitor (100 nM and 300 nM). Significant reduction in formation of neurite outgrowths was observed in TGs when cultured with different concentrations of inhibitor compared to control group (Fig.S3J,L) revealing the important role of both CDK13 and CDK12 in development of peripheral nerves.

### Changes in expression of key developmental molecules in early craniofacial development of *Cdk13*-deficient embryos

We also detected alterations in expression patterns of genes and proteins strongly associated with craniofacial development and specifically enriched in mesenchymal (SOX9, *Msx1*, *Foxd1*) and epithelial structures (SOX2, *Meis2*, *Shh*) in this region. In E11.5 *Cdk13^tm1d/tm1d^*embryos, we observed almost absence of neural crest marker SOX9 in the mesenchyme of lateral nasal prominences (lnp) and maxillary prominences (mx), including area with undeveloped MxN (Fig.4H′). On the other hand, pattern of SOX2, one of regulators of neural cells differentiation and peripheral nervous system development (Adameyko et al., 2012), was similar to WT embryos, only missing in the area of the prospective MxN (Fig.4Í). *Meis2*, a gene involved in cranial neural crest development (Machon et al., 2015), was absent from the neural tube and enriched in the mesenchyme of medial nasal prominences (Fig.4D) and *Msx1*, a key player in craniofacial development (Levi et al., 2006), was detected absent from mesenchyme of the medial nasal prominences (Fig.4E). Detection of genes necessary for determination and development of facial primordia (Jeong et al., 2004), showed enhanced expression of *Shh* in the ventral neural tube and its altered pattern in the epithelium covering forming stomodeal cavity (Fig.4F) and enhanced expression of *Shh* downstream gene *Foxd1* in the developing mandibular prominences (Fig.4G). It confirms that *Cdk13*-deficiency leads to deregulation of the key developmental genes already in early craniofacial development which subsequently results in severe facial clefting.

### *Cdk13*-deficiency alters expression of neurogenesis-specific genes and key morphogenic molecules in the developing secondary palate

To further evaluate alterations in gene expression of molecules related to neurogenesis during palatogenesis in *Cdk13^tm1a/tm1a^* embryos, we used PCR Arrays where 84 genes could be analyzed simultaneously. We observed differences in expression of several genes in the rostral and caudal areas of the palatal shelves at both selected stages (E12.5 and E14.5) as response to *Cdk13*-deficiency (Fig.5A-D). Significant changes were found in molecules involved in apoptosis (*Adora2a, S100b, Vegfa*), neuronal differentiation (*Cdk5rap2, Egf, Hes1, Mef2c, Neurog1, Noggin, Olig2, S100b*), cellular adhesion (*Nrcam, Tnr*), neural migration (*Ndn*), axonogenesis (*Egf, Nrcam, S100a6, S100b, Tnr*), synaptogenesis (*Ache*), synaptic transmission (*Chrm2, Drd2, Sod1*), and regulation of production of cytokines and growth factors (*Cxcl1, Gpi1, Mdk, Ptn, S100a6, Tgfb1, Vegfa*) in *Cdk13*-deficient animals.

**Figure 5.**
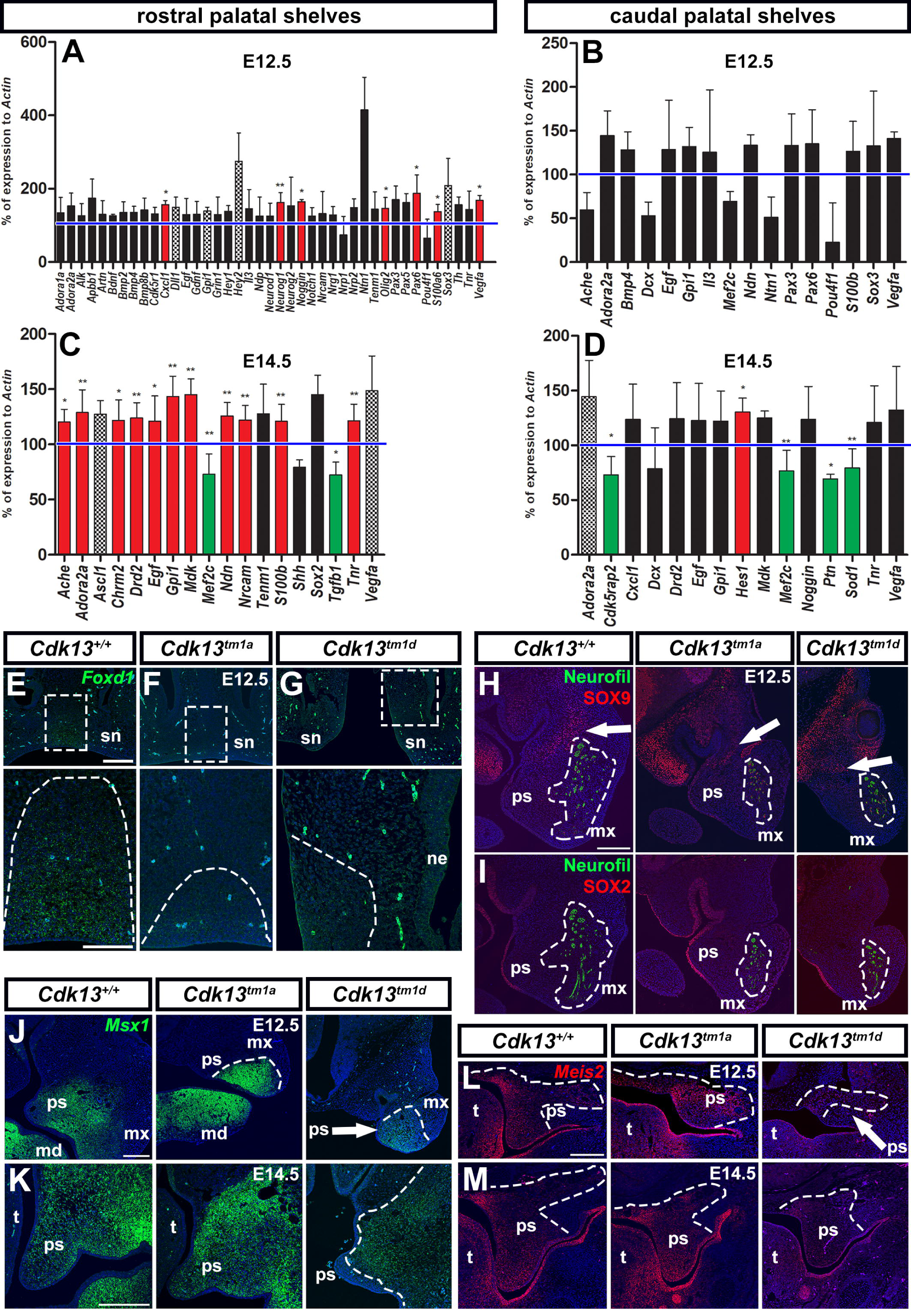
Gene expression quantification analysis of the neurogenesis-associated molecules in the *Cdk13^tm1a/tm1a^* palatal shelves and *in situ* gene and protein expression on sections. **(A-D)** PCR Array for mouse neurogenesis specific genes performed separately on tissues dissected from the E12.5 and E14.5 rostral and caudal palatal shelves of *Cdk13^tm1a/tm1a^*embryos. (A) In the E12.5 rostral palatal shelves, significantly increased gene expression was observed in *Cxcl1* (*C-X-C chemokine ligand 1*), *Neurog1* (*Neurogenin 1*), *Nog* (*Noggin*), *Olig2* (*Oligodendrocyte transcription factor 2*), *Pax6* (*Paired box gene 6*), *S100a6* (*Calcyclin*) and *Vegfa* (*Vascular endothelial growth factor A*). (B) In the E12.5 caudal palatal shelves, expression of 16 genes was altered but no statistically significant change in the level of expression was detected. (C) In the E14.5 rostral palatal shelves, *Ache* (*Acetylcholinesterase*), *Adora2a* (*Adenosine A2a receptor*), *Chrm2* (*Cholinergic receptor, muscarinic 2*), *Drd2* (*Dopamine receptor D2*), *Egf* (*Epidermal growth factor*), *Gpi1 (Glucose phosphate isomerase 1)*, *Mdk* (*Midkine*), *Ndn* (*Necdin*), *Nrcam* (*Neuronal cell adhesion molecule*), *S100b* (*S100 Calcium-Binding Protein B*) and *Tnr* (*Tenascin R*) expression were significantly increased, while *Mef2c* (*Myocyte enhancer factor 2c*) and *Tgfb1* (*Transforming growth factor, beta 1*) gene expression was significantly decreased. (D) In the E14.5 caudal palatal shelves, significantly increased expression was observed in *Hes1* (*Hairy and enhancer of split 1*), while significantly decreased gene expression was detected in *Cdk5rap2* (*cyclin-dependent kinase 5 regulatory subunit associated protein 2*), *Mef2c*, *Ptn* (*Pleiotrophin*) and *Sod1* (*Superoxide dismutase 1*). Red columns represent significantly increased gene expression, green columns significantly decreased gene expression, patterned columns display marginally significant changes in gene expression and black columns not significantly changed expression. Level of changed expression was set up at 120 % for fold up, and 80 % for fold down changes in % of expression to *Actin* expression. Blue lines highlight 100 % level. Unpaired two-tailed Student t - test; **0.001 < p < 0.01; *p < 0.05; 0.05 < p < 0.1 (marginally significant). **(E-G)** RNAScope detection of *Foxd1* (green) *in situ* gene expression on sections of E12.5 embryos from the rostral snout region. White dashed line rectangles highlight regions for detailed pictures. Dashed lines in detailed pictures outline *Foxd1* expression domains **(H-I)** Immunohistochemical detection of neurofilaments (green, anti-2H3), SOX9 (red) and SOX2 (red) expression on section of E12.5 embryos. White arrows point to regions enriched with SOX9-positive cells. White dashed lines outline regions with 2H3 signal representing developing maxillary nerves. **(J-K)** RNAScope detection of *Msx1* (green) *in situ* gene expression on sections of E12.5 and E14.5 embryos from the rostral palate region. White dashed lines outline *Msx1-*positive regions. **(L-M)** RNAScope detection of *Meis2* (red) *in situ* gene expression on sections of E12.5 and E14.5 embryos from the caudal palate region. White dashed lines outline *Meis2-*positive regions. Scale bars: 200 μm; *Foxd1* detail – 100 μm. md – mandibular prominence; mx – maxillary prominence; ne – nasal epithelium; ps – palatal shelves; sn – snout; t – tongue.

Moreover, altered expression of palatal genes and proteins was detected later at E12.5 or E14.5 respectively. *Cdk13*-deficiency lead to noticeable reduction of *Foxd1* in the mesenchyme of snout midline (Fig.5F,G), *Msx1* reduction in the rostral palatal shelves and also reduction of *Meis2* in the caudal palatal shelves (Fig.5J,K), especially at E12.5. Expression of both SOX9 and SOX2 was similar to pattern observed in wt animals (Fig.5H,I).

## DISCUSSION

### CDK13 controls craniofacial morphogenesis through regulation of expression of responsible developmental pathways

Here, we determined a key contribution of CDK13 to the craniofacial morphogenesis and neurogenesis, where the phenotype of *Cdk13-*deficient animals included smaller head and disturbed facial morphology, including facial clefts. In our mutants, we found altered expression of *Msx1*, *Meis2*, *Shh* and *Foxd1* in craniofacial structures. Downregulation or upregulation of these genes or their downstream targets causes similar phenotypic craniofacial phenotype including facial clefts or reduced growth of facial nerves (Levi et al., 2006, Machon et al., 2015, Jeong et al., 2004). All the affected craniofacial structures are mostly of NCCs origin, which marker SOX9 was also reduced in *Cdk13*-deficient animals and moreover, we detected significant downregulation of *Mef2c*, a direct transcriptional target of SOX10 (important transcription factor for development of NCCs derivatives), (Agarwal et al., 2011) which conditional mutation in NCCs results in craniofacial defects and neonatal lethality (Verzi et al., 2007).

### Defects in craniofacial morphogenesis and neurogenesis as a result of disrupted CDK-associated signaling

Similarly to patients with mutated *CDK13*, mutations in genes encoding other CDKs and their associated proteins, such as cyclin M, cyclin K, CDK10 (Colas, 2020), and CDK5RAP2 (Yigit et al., 2015) produce diverse phenotypes. These patients exhibited not just craniofacial defects but also affected neural tissues: mutation in *cyclin M* resulted in optic nerve hypoplasia, mutation in *cyclin K* led to developmental delay and intellectual disabilities and mutations in *CDK10* caused intellectual disabilities connected with language and learning disorders. Patients with loss-of-function mutation of *CDK5* displayed craniofacial defects other than facial clefts, (short forehead, full cheeks, micrognathia), they also suffer with lissencephaly, cerebellar hypoplasia, microcephaly, intellectual disabilities, speech delay or autistic features (Colas, 2020), similar to patients with *CDK13* mutations (Hamilton and Suri, 2019).

CDK5 has specific role in neural tissues physiology and regulates several neural processes such as an axonal transport, migration, synaptic vesicle endocytosis and is also important for regulation of axon and neurite outgrowth (Shah and Rossie, 2018). Its knockout leads to perinatal lethality in mice connected with deficient neuronal migration and impaired axonal transport of neurofilaments (Ohshima et al., 1996). Importantly, i*n vitro* experiments in cortical neurons demonstrated the role of the CDK13 and also CDK12 in regulation of neurite outgrowth through a common signaling pathway that involves modulation of *Cdk5* at the RNA level (Chen et al. 2014). Here we found significant downregulation of one of the CDK5 regulatory proteins, *Cdk5rap2*, in the E14.5 caudal palate, and conversely upregulated (not significantly) *Cdk5r1* regulatory subunit in the E12.5 rostral palate confirming CDK13-CDK5 functional association observed in *in vitro* experiments. Moreover, *CDK5RAP2* mutations in human patients cause Seckel syndrome manifesting by microcephaly and cognitive problems (Yigit et al., 2015) and *CDK5R1* was proposed as a candidate gene in patients affected by NF1 microdeletion syndrome which mutations lead to non-syndromic intellectual disability and undefined facial deformities (Venturin et al., 2006). Therefore, CDK5-signaling can be probably regulated by CDK13 activity during facial morphogenesis and neurogenesis, as suggested by its localization in growing neurites of cultured neurons, which however will be necessary to follow in future.

### Defective peripheral nerves development as a potential cause of craniofacial defects

Our findings uncovered that *Cdk13*-deficiency triggers the alteration of neurogenesis-specific genes in the PSs and development of hypoplastic craniofacial nerves *in vivo* confirming previous *in vitro* study in cortical neurons (Chen et al., 2014). We hypothesized, that defective development of peripheral nerves and associated cells in *Cdk13*-deficient animals can lead to secondary phenotype in form of facial defects. Such non-canonical function of nerves as moderators of facial morphogenesis has been proven in Möbius syndrome, where deficient innervation by abducens and facial nerves lead to cleft palate, tongue and midfacial hypoplasia (Rizos et al., 1998) associated with mutations in *TUBB3* (a gene encoding tubulins expressed in neurons important for axon guidance) (Patel et al., 2017) or *PLXND1* (a gene encoding plexin cell surface receptor for semaphorins located in lamellipodium and important for cell migration) (Tomas-Roca et al., 2015). On the contrary, based on other studies, Möbius syndrome is suggested to be caused by developmental defects of the entire rhombencephalon rather than just underdevelopment of craniofacial nerves (Verzijl et al., 2003). Another disorder of peripheral nervous system which leads to cleft palate and other facial defects, is known as hereditary sensory and autonomic neuropathy type IV (HSAN IV) (Gao et al., 2013). This disorder is associated with mutation in *NTRK1*, a gene encoding tyrosine kinase receptor (TRKA) for nerve growth factor (NGF) which is important also for craniofacial development (Louryan et al., 1995) and its mutation led to reduced neurite growth *in vitro* (Nakajima et al., 2020). Hemifacial atrophy was seen in Parry-Romberg syndrome accompanied by various neurological pathologies, unfortunately with unknown etiology (Vix et al., 2015).

CDK13 is ubiquitously expressed in murine snout, developing PSs and craniofacial nerves and its mutation leads to prominent facial clefts. Craniofacial structures, including palate, develop with contribution of the neural crest cells (NCCs). After delamination from neural tube, NCCs migrate either by free migration or along the peripheral nerves based on developmental stage. Growing peripheral nerves and glial cell types originate mostly from migrating NCCs and Schwann cell precursors (SCPs) (Furlan and Adameyko, 2018), but there are few exceptions, among them sensory neurons which innervate maxillary region including palate. These neurons originate in mesencephalic trigeminal nucleus in CNS and their development is regulated by FGF8 and Pou4f1 (*Pou4f1* is downregulated in *Cdk13*-deficient PSs), (Hunter et al., 2001). We detected changes in expression of genes specific for migration and axon outgrowth, such as *Egf* which induces axon growth (Onesto et al., 2021) as well as changes in expression of *Nrg1* which encodes EGF ligand, expressed in trigeminal ganglion (Meyer et al., 1997). Interestingly, *Nrg1*-deficiency in mice leads to maxillary dysmorphology (Waddington et al., 2017). We also detected upregulation of other genes known to be expressed in the trigeminal ganglion: *Neurog1* also expressed in PSs (Visel et al., 2007) and *Nrp2* encoding receptor for SEMA3C/3F, axon guidance factors (Chen et al., 1997) also expressed in migrating NCCs (Maden et al., 2012). Similarly to our findings in PSs, *Neurog1* and *Nrp2* were upregulated in trigeminal ganglion as a result of *Pou4f1* downregulation (Lanier et al., 2009). This deregulated system exhibits tendency to rescue growth of nerves in the rostral palate as suggested by extreme upregulation of *Ntn1*, a gene encoding secreted factor responsible for guidance of developing peripheral motor axons (Serafini et al., 1996) and which was proposed to be one of the risk factors in development of the non-syndromic cleft lip and cleft palate (Leslie et al., 2017).

The formation of myelin sheet from glial cells around nerves is important for their proper development and function provided by Schwan cells (SCs). We detected upregulated *S100b*, a gene specifically expressed in myelinating and non-myelinating SCs and also in satellite glial cells (Avraham et al., 2022). Differentiation of SCs and axon myelination is strongly dependent on Notch signaling (Woodhoo et al., 2009). Interestingly, here we showed that *Cdk13^tm1a/tm1a^* mutation led to the upregulation, although nonsignificant, of several Notch-related genes (*Ascl1*, *Notch1*, *Dll1*, *Hey1/2)* and later in development, to upregulation of Notch downstream target *Hes1,* which is important for development of craniofacial region including palate (Akimoto et al., 2010) and cranial and spinal peripheral nerves (Hatakeyama et al., 2006).

In *Cdk13*-deficient animals, we also determined alterations in gene expression of molecules involved in adhesion and migration which could lead to altered nerve growth. *Nrcam* encodes adhesive molecule involved in neurite outgrowth, axonal pathfinding or cell migration and its mutation in human lead to developmental delay or peripheral neuropathy (Kurolap et al., 2022). Upregulation of *Nrcam* was associated with cleft palate caused by defective TGF-β signaling (Liu et al., 2020). Similarly, we detected significantly upregulated *Nrcam* in the caudal palate and significantly downregulated *Tgfb1*, other member of TGF-β signaling.

Important molecules for the migration, axon outgrowth and survival of developing nerves or neural progenitors are chemokines or neurotrophic factors. We detected changes in expression of such factors, among them *Gpi1* (Haga et al., 2000) and *Vegfa* (McLennan et al., 2010) which were upregulated in all the analyzed regions. This can be explained by the attempt of mutants to attract more migrating cells and growing axons as response to *Cdk13*-deficiency. It is also supported by upregulation of *Egf* in all the palatal regions we analyzed. Moreover, we revealed upregulation of *Adora2a,* which positively regulates expression of neurotransmitter acetylcholine and when mutated it increases autistic symptoms and anxiety in human (Freitag et al., 2010) similarly to patients with mutation in *CDK13* (Hamilton and Suri, 2019, Rouxel et al., 2022). In addition, we detected changes in gene expression of other synapse-associated genes encoding namely *Ache*, *Chrm2*, *Drd2* and *Sod1* which could lead to altered synaptogenesis and synaptic transmission in our mutants.

All these gene expression changes together with hypoplastic craniofacial nerves and ganglia suggest the role of CDK13 in neurogenesis during craniofacial development probably through affecting cellular migration and neural outgrowth. Based on our findings, we propose functional and structural role of CDK13 in formation of cellular outgrowths, especially long neural outgrowths, based on its localization to cellular protrusions. Its deficiency thus could lead to altered intracellular transport of important cytoskeletal, adhesion, migration and synaptic components, which are critical especially for formation of long protrusions.

Nevertheless, functional association between development of hypoplastic peripheral craniofacial nerves which would lead secondary to development of craniofacial clefts in *Cdk13*-deficient embryos, is missing. To prove this association, it will be necessary to generate conditional mouse mutants with either ablation of specific cell population or more generally, ablate the trigeminal nerve branches to block possible non-canonical function of these nerves in facial morphogenesis.

### CONCLUSIONS

CDK13 is a key molecule for development of numerous tissues and organs including craniofacial structures. Here, we determined its role during development of facial structures where loss-of-function in *Cdk13* results in cleft lip/palate and formation of midfacial cleft. *Cdk13*-deficient animals exhibit altered neurogenesis caused by distorted expression of neurotrophic molecules leading to development of hypoplastic craniofacial nerves.

## MATERIALS AND METHODS

### Embryonic material

Heterozygous hypomorphic (*Cdk13^tm1a(EUCOM)Hmgu^*) and knock out (*Cdk13^tm1d^*) animals were obtained from the Infrafrontier Research Infrastructure – Mouse disease Models. Mouse were generated at the Transgenic and Archiving Module, CCP (IMG, Prague, Czech Republic). Breeding and genotyping protocols were performed as previously described elsewhere (Nováková et al., 2019). All animal procedures were performed in strict accordance with the Guide for the Care and Use of Laboratory Animals and approved by the Institutional Animal Care and Use Committee (Masaryk University, Brno, Czechia, No. MSMT-34505/2020-7).

### Scanning Electron Microscopy

Embryos (control and *Cdk13^tm1a/tm1a^*) were fixed in 4% paraformaldehyde, washed in distilled water and dehydrated through a graded series (30-100%) of ethanol solutions. Later, samples were dried out using CPD 030 Critical Point Dryer (BAL-TEC) and shadowed by gold in metal shadowing apparatus Balzers SCD040. Samples were observed and photographed in the scanning electron microscope TESCAN Vega TS 5136 XM (Tescan, Czech Republic). SEM was performed on 1 embryo of each stage (E12.5, E13.5, E14.5, E16.5) with representative phenotype.

### Immunofluorescence on slides

Embryonic tissues were fixed in 4% PFA overnight. Specimens were then embedded in paraffin and cut in transversal and sagittal planes for 5 µm sections. For immunohistochemistry staining, sections were deparaffinized in xylen and rehydrated in an ethanol series. Antigen retrieval was performed either in 1% citrate buffer or in DAKO antigen retrieval solution (S1699, DAKO Agilent, USA) at 97.5 °C.

For protein localization, we incubated sections with primary antibody for 1 hour at room temperature or overnight at 4 °C. Following antibodies were used: 2H3 (1:50, Nefm, AB_2314897, Developmental Studies Hybridoma Bank), SOX2 (1:100, 2748s, Cell Signaling), SOX9 (1:100, HPA001758, Sigma), Ki67 (1:200, RBK027, Zytomed systems). Then sections were incubated with following secondary antibodies (1:200) for 30 minutes at room temperature: anti-mouse Alexa Fluor 488 (A11001), and anti-rabbit Alexa Fluor 594 (A11037, both Thermo Fisher Scientific, USA). Ki-67 positive cells were detected by anti-rabbit secondary antibody, ABC binding (PK-6101, Vector laboratories) and followed by application of the DAB (K3468, Dako) chromogenic system.

Sections were mounted with Fluoroshield with DAPI (F6057, Sigma, Merck, Germany). If DR (62251, Thermo Fisher Scientific, USA) was used for nuclei staining, sections were mounted with Fluoroshield (F6182, Sigma, Merck, Germany). Pictures were taken on confocal microscopes Leica SP8 (Leica, Germany) and Zeiss LSM800 (Zeiss, Germany). Nuclei on DAB stained sections were counterstained with hematoxilyn. Sections were photographed under bright-field illumination with the Leica DMLB2 compound microscope (Leica, Germany).

Mitotic index on Ki-67 stained sections in the palatal shelves, was counted as a ratio between Ki-67-positive cells (brown) and total number of cells (negative plus Ki-67-positive cells). Cells were counted independently in mesenchyme and epithelium of the palatal shelves. Cells were counted on 4 sections (both left and right palatal shelves) in 3 embryos for each genotype (*Cdk13^+/+^* and *Cdk13^tm1a/tm1a^*).

### Immunocytochemistry on glass inserts

Mesenchymal embryonic fibroblasts (MEF) and cells from embryonic DRGs (dorsal root ganglia) were isolated from E12.5 embryonic bodies. Tissues were enzymatically processed using Dispase II (D4693, Sigma, Merck, Germany) for 1 hour at 37 °C while shaking. Cells were then centrifuged, filtered through 40 µm Cell strainer (431750, Corning), seeded on glass inserts and left to grow until 70 – 80 % confluency in DMEM cultivation medium (D6546, Sigma, Merck, Germany). Adult DRG cells were isolated from DRGs from adult mice. DRGs were enzymatically processed using Collagenase IV (LS0004188, PAN Biotech) for 6 hours at 37 °C. Tissues were resuspended every hour by pipetting. Cells were then filtered through Cell strainer and seeded on glass inserts and left to grow and form long outgrowths in Neurobasal cultivation medium (21103-49, Gibco). Cells were then fixed in 4% PFA for 15 minutes.

For protein localization, we incubated cells on glass inserts with primary antibody for 1 hour at room temperature. Following antibodies were used: anti-CDK13 (1:100, HPA059241, Sigma, Merck, Germany), anti-CDK13 N-TERM (1:100, SAB1302350, Sigma), anti-F-actin (1:100, A12379, Alexa Fluor 488™ phalloidin, Thermo Fisher), anti-Sodium Potassium ATPase (1:100, ab76020, Abcam), anti-2H3 (1:50, Nefm, AB_2314897, Developmental Studies Hybridoma Bank). Then sections were incubated with following secondary antibodies (1:200) for 30 minutes at room temperature: anti-mouse Alexa Fluor 488 (A11001), and anti-rabbit Alexa Fluor 594 (A11037, both Thermo Fisher Scientific, USA). Glass inserts were mounted on glass slides with Fluoroshield with DAPI (F6057, Sigma, Merck, Germany). Pictures were taken on confocal microscope Leica SP8 (Leica, Germany). Cells were cultivated from at least 3 different embryos of all three genotypes (*Cdk13^+/+^*, *Cdk13^tm1a/tm1a^*and *Cdk13^tm1d/tm1d^*).

### Whole mount immunofluorescence

Embryos were dissected, fixed in 4% PFA while rotating at 4 °C for 4 hours. Embryos were then postfixed in methanol through gradual concentrations of methanol (25%, 50%, 75%, 100%). Embryos were then bleached in mixture of hydrogen peroxide, DMSO and methanol for 24 hours and then postfixed in combination of DMSO and methanol. Embryos were incubated with primary antibody to neurofilaments (2H3, AB_2314897, Developmental Studies Hybridoma Bank) for 7 days while rotating and then with secondary antibody (anti-mouse Alexa Fluor 488, A11001) for 2 days while rotating. Embryos were finally cleared in mixture of benzyl benzoate and benzyl alcohol until they get transparent. For microscopy, we placed embryos on NuncTM Glass Bottom dish (150680, Thermo Fisher) and imaged them on Zeiss AxioZoom.V16-Apotome2 (Zeiss, Germany) at CELLIM (Core Facility Cellular Imaging, CEITEC, Masaryk University, Brno, Czech Republic). Whole mount immunodetection of the neurofilaments was performed in 3 different embryos of all three genotypes (*Cdk13^+/+^*, *Cdk13^tm1a/tm1a^* and *Cdk13^tm1d/tm1d^*) at E11.5 stage.

### PCR arrays analysis

PCR arrays were performed on tissues isolated from rostral and caudal parts of the palatal shelves from E12.5 and E14.5 embryos. One sample was pooled from two or three embryos, three biological replicates for each stage and genotype were analyzed. Total RNA was extracted using RNeasy Plus Mini Kit (74136, Qiagen, Germany) according to the manufacturer’s instructions. Total RNA concentration and purity was measured using a NanoDrop One (Thermo Fisher Scientific, USA). First-strand cDNA was synthesized using gb Reverse Transcription Kit (3012, Generi Biotech, Czech Republic) according to the manufacturer’s instructions. RT² Profiler™ PCR Array Mouse Neurogenesis (330231, Qiagen, Germany) was performed according to the manufacturer’s instructions on LightCycler 96 (Roche, Germany). Data were evaluated by ΔΔCT method with normalization against *Actin* levels.

### RNAScope

The embryos were fixed in 4% PFA and fixation time differed based on the stage. The tissues were then dehydrated in an ethanol series, embedded in paraffin, and 5 µm transverse sections were obtained. The sections were deparaffinized in xylene and dehydrated in 100% ethanol. To detect gene expression, we used the RNAScope Multiplex Fluorescent v2 Assay kit (323 110, ACD Bio, United States) for formalin-fixed paraffin embedded tissues according to the manufacturer‘s instructions. All reactions, which require 40 °C incubation temperature, were performed in the HybEZTM II Oven (ACD Bio, United States). Probes for *Cdk13* (895581), *FoxD1* (495501), *Meis2* (436371), *Msx1* (421841) and *Shh* (314361), all ACD Bio,

United States, were used. The hybridized probes were visualized using the TSA-Plus Cyanine 3 system (NEL744001KT, Perkin-Elmer, United States), according to the manufacturer’s protocol. DAPI (323 108, ACD Bio, United States) was used to stain nuclei. Pictures were obtained with the Leica SP8 confocal microscope (Leica, Germany).

### RT-PCR

Rostral and caudal parts of the palatal shelves (E12.5, E14.5, E16.5), and maxillary (mx), mandibular (md) and frontonasal prominences (fnp) (E11.5, E12.5) from WT embryos were dissected to quantify differences in *Cdk13* expression during development of the facial structures. Individual parts of the palatal shelves and facial prominences were dissected from at least 3 different embryos for each stage. Proliferation rate changes in the *Cdk13*-deficient embryos were assessed using *CyclinD1* gene expression in tissues isolated from the rostral and caudal palatal shelves (*Cdk13^tm1a^* E12.5, E14.5) and from lips (*Cdk13^tm1d^* E12.5). Total RNA was extracted using RNeasy Plus Mini Kit (74136, Qiagen, Germany) according to the manufacturer’s instructions. Total RNA concentration and purity was measured using a NanoDrop One (Thermo Fisher Scientific, USA). First-strand cDNA was synthesized using gb Reverse Transcription Kit (3012, Generi Biotech, Czech Republic) according to the manufacturer’s instructions. TaqMan probe was used to quantify *Cdk13* (Mm01164725_m1, Thermo Fisher) and *CyclinD1* (Mm00432359_m1, Thermo Fisher) gene expression. The RT-PCR reaction was performed on LightCycler 96 (Roche, Germany). The comparative C_T_ method was used for analysis.

### Trigeminal ganglia cultivation and neurite outgrowth assay

Trigeminal ganglia were dissected from E12 embryos and washed in ice cold Neurobasal medium. Each TG was placed in a separate well on culture well plate into 10 μl drop of Matrigel (356231, Corning, USA) and left 20 minutes to polymerize in the tissue culture incubator. Neurobasal medium (21103049, Gibco, Thermo Fisher, USA) supplemented with 100 nM or 300 nM THZ531 inhibitor (SML2619, Sigma Aldrich, Merck, Germany) was added. Pictures were snapped after adding cultivation medium (0 h) and then after 24 hours using Olympus IX71 inverted microscope (Olympus, Japan). Neurite outgrowth was measured using ImageJ software (NIH, USA) while comparing areas covered by neurites on pictures of different culture conditions. Percentage changes in ability of TGs to produce neurite outgrowths were calculated as following: *neurite outgrowth area (total area – TG area)/total area*100*.

### Terminal Deoxynucleotidyl Transferase dUTP Nick End Labeling (TUNEL) Assay

Apoptotic cells were detected using the TUNEL assay (ApopTag Peroxidase In Situ Apoptosis Detection Kit, Cat. No. S7101, Chemicon, Temecula, United States). Nuclei were counterstained with hematoxylin. Sections were photographed under bright-field illumination with the Leica DMLB2 compound microscope (Leica, Germany). TUNEL Assay was performed in 3 different embryos of *Cdk13^+/+^* and *Cdk13^tm1a/tm1a^* genotypes.

### Statistical analyses

Data were evaluated for statistical significance in GraphPad (GraphPad Software, Boston, MA, USA) using unpaired two-tailed Student’s t-tests. Differences were considered to be significant at p<0.05, p<0.01 and p<0.001 and were indicated by *, ** or *** symbols, respectively.

## Acknowledgement

This work was supported by the Czech Science Foundation (19-01205S) and by the MEYS CR (CZ.02.1.01/0.0/0.0/15_003/0000460). We acknowledge the core facility CELLIM supported by the Czech-BioImaging large RI project (LM2018129 funded by MEYS CR).

## Author contributions

M. Hampl, contributed to conceptualization and design, data acquisition, analysis, visualization and interpretation, drafted the manuscript; N. Jandova, contributed to data acquisition, analysis, visualization and interpretation, critically revised the manuscript; D. Luskova, contributed by data curation, resources, methodology; M. Novakova, contributed to data acquisition, analysis and interpretation, J. Prochazka, J. Kohoutek, M. Buchtova, contributed to conception and design, critically revised the manuscript. All authors gave final approval and agreed to be accountable for all aspects of the work.

